# Testing the efficacy of different molecular tools for parasite conservation genetics: a case study using horsehair worms (Phylum Nematomorpha)

**DOI:** 10.1101/2023.02.21.529467

**Authors:** Mattia De Vivo, Wei-Yun Chen, Jen-Pan Huang

**Affiliations:** Department of Life Science, National Taiwan Normal University, Taipei, Taiwan; Biodiversity Program, Taiwan International Graduate Program, Taipei, Taiwan; Biodiversity Research Center, Academia Sinica, Taipei, Taiwan

**Keywords:** Parasite conservation, Nematomorpha, Taiwan, COXI, ddRADseq, population size

## Abstract

In recent years, parasite conservation has become a globally significant issue. Because of this, there is a need for standardised methods for inferring population status and possible cryptic diversity. However, given the lack of molecular data for some groups, it is challenging to establish procedures for genetic diversity estimation. Therefore, universal tools, such as double digest restriction-site associated DNA sequencing (ddRADseq), could be useful when conducting conservation genetic studies on rarely studied parasites. Here, we generated a ddRADseq dataset that includes all three described Taiwanese horsehair worms (Phylum Nematomorpha), possibly one of the most understudied animal groups. Additionally, we produced data for a fragment of the cytochrome c oxidase subunit I (COXI) for said species. We used the COXI dataset in combination with previously published sequences of the same locus for inferring the effective population size (*Ne*) trends and possible population structure.

We found that a larger and geographically broader sample size combined with more sequenced loci resulted in a better estimation of changes in *Ne*. We were able to detect demographic changes associated with Pleistocene events in all the species. Furthermore, the ddRADseq dataset for *Chordodes formosanus* did not reveal a genetic structure based on geography, implying a great dispersal ability, possibly due to its hosts. We showed that different molecular tools can be used to reveal genetic structure and demographic history at different historical times and geographical scales, which can help with conservation genetic studies in rarely studied parasites.

## Introduction

In recent years, there has been a growing concern regarding the conservation of parasites (e.g., Koh *et al.*, 2004; Dobson *et al.*, 2008; Dunn *et al.*, 2009; Gómez and Nichols, 2013; Dougherty *et al.*, 2016; Carlson *et al.*, 2017, 2020; Kwak *et al.*, 2020). This is because parasites are integral parts of the ecosystems from several points of view, i.e., species diversity (Poulin and Morand, 2004), biomass (Kuris *et al.*, 2008) and their importance in food webs (Lafferty *et al.*, 2008; Sato *et al.*, 2008, 2011, 2012; Amundsen *et al.*, 2009; Dougherty *et al.*, 2016). Moreover, they suffer from human-made environmental changes and can go extinct together with their hosts (Dunn *et al.*, 2009; Carlson *et al.*, 2017, 2020). However, given that parasites have only recently started to be regarded as important from a conservation standpoint, tools for parasite conservation have rarely been tested (Carlson *et al.*, 2020; Poulin, 2021). Particularly, molecular tools have not been extensively used for parasites (Criscione *et al.*, 2005; Criscione, 2016; Selbach *et al.*, 2019). These tools can be important for inferring a population’s cryptic genetic diversity and changes in effective population size (*Ne*) over time (Criscione, 2013, 2016; Strobel *et al.*, 2019). *Ne* is known to greatly influence the genetic diversity and viability of populations (Criscione *et al.*, 2005; Franklin, 1980; Pérez□Pereira *et al.*, 2022), but estimating it in parasites may come with some caveats depending on the studied system (Criscione and Blouin, 2005; Strobel *et al.*, 2019). That being said, estimates of *Ne* can be helpful for considering the taxa’s conservation status (Franklin, 1980; Pérez□Pereira *et al.*, 2022). Unfortunately, the current lack of data makes it difficult for researchers to enact timely conservation plans, putting some species at risk of extinction (Carlson *et al.*, 2017, 2020; Poulin, 2021).

Methods for estimating *Ne* trends from genetic data of non-model organisms are being developed (e.g., Liu and Fu, 2020). This has been possible thanks to protocols that allow researchers to generate genome-wide data without reference sequences. For example, restriction-site associated DNA sequencing (RADseq) approaches have been shown to work well with non-model organisms and became popular in population and conservation genetic studies thanks to their relatively cheap cost and versatility (Miller *et al.*, 2007; Andrews *et al.*, 2016; Marandel *et al.*, 2020). This is interesting from a molecular conservation parasitology perspective, given that non-medically relevant parasites often lack reference data (Selbach *et al.*, 2019). Alternatively, even single-locus methods for *Ne* estimation are available, allowing researchers to use previously released data (i.e., sequences from GenBank) for demographic inferences, although having more loci leads to better parameter estimation (Vitalis and Couvet, 2001; Ho and Shapiro, 2011). Single-locus or double-locus data are available for some parasites, and it is expected that their number in public databases will increase given their use in DNA barcoding (Selbach *et al*., 2019).

Among parasitic taxa with relatively low data availability, horsehair worms (phylum Nematomorpha) have interesting characteristics from both conservation and ecological perspectives. All the taxa in this phylum are parasites with a free-living adult stage (Schmidt-Rhaesa, 2012; Bolek *et al.*, 2015), making Nematomorpha one of the few phyla in which all the known species are parasitic (see Giribet and Edgecombe, 2020). The most specious group, which includes worms called “gordiids” (Gordioida; but see Schmidt-Rhaesa 2012), is notoriously known for manipulating their final hosts to jump into water, where the adult worms are released (Thomas *et al.*, 2002; but see Valvassori *et al.* 1988, Schmidt-Rhaesa and Kristensen 2006, Chiu *et al*. 2020 and Anaya *et al*. 2021 for alternative lifecycles). Such manipulation seems to be achieved by making the hosts more sensitive to polarized light (Obayashi *et al*., 2021). This led to the exaggerated claim that these worms can totally control their final hosts’ movements and make them commit suicide, giving gordiids a misleading reputation and a popular science notoriety at the same time (Doherty, 2020). Nevertheless, it is evident that they can impact food networks in a community by making their hosts jump into water, which makes said hosts easy prey to their predators (Sato *et al.*, 2008, 2011, 2012). Although common in suitable environments (Schmidt-Rhaesa, 2012; Chiu, 2017), it is known that pollution and human-made changes (i.e., clear-cut logging and stream remediation) can have negative effects on hairworms, even causing local extinction in some cases (Poinar, 2008; Sato *et al.*, 2014; Chiu *et al.*, 2016; Achiorno *et al.*, 2018).

In this study, multiple protocols and methods were used for generating molecular data to study population structure and the changes of *Ne* over time in gordiids; the same methods can be applied to other parasitic groups for conservation and biodiversity-oriented studies. Specifically, a double digest restriction-site associated DNA sequencing (ddRADseq) dataset including all three known horsehair species in Taiwan was generated. Additionally, DNA sequence data from the mitochondrial cytochrome c oxidase subunit I (COXI) region were generated and used for evaluating population and genetic status in combination with previously released data (Chiu *et al.*, 2011, 2016, 2017, 2020). With both datasets, population structure and demographic history were estimated and compared. More focus was given to *Chordodes formosanus*, since they are relatively easier to sample and are more common (Chiu, 2017). A nation-wide ddRADseq was generated for said species, allowing us to estimate *Ne* and population structure at a nation-wide level. Additionally, the Taiwanese and Japanese populations’ trends were compared using the COXI dataset.

## Materials and methods

### Sampling

Gordiids were sampled over a span of 14 years, from 2007 to 2021 (Fig. 1: Sup. Table 1). *Chordodes formosanus*, which is regarded as the most common species in Taiwan (Chiu, 2017), was sampled nationwide, while specimens of *Acutogordius taiwanensis* and *Gordius chiashanus* were collected from fewer localities (Fig. 1; Sup. Table 1). Specimens were collected either by hand in bodies of water (i.e., streams and man-made ponds), by sampling the hosts and then immersing them until the adult worm came out, or by collecting the dead gordiids on the ground, which is the most common method for finding them in Taiwan (Chiu, 2017). The samples were fixed at 95% alcohol and stored at −20°C. Species recognition was based on DNA barcoding, given that the majority of the samples were too ruined for morphological identification.

**Figure 1.**
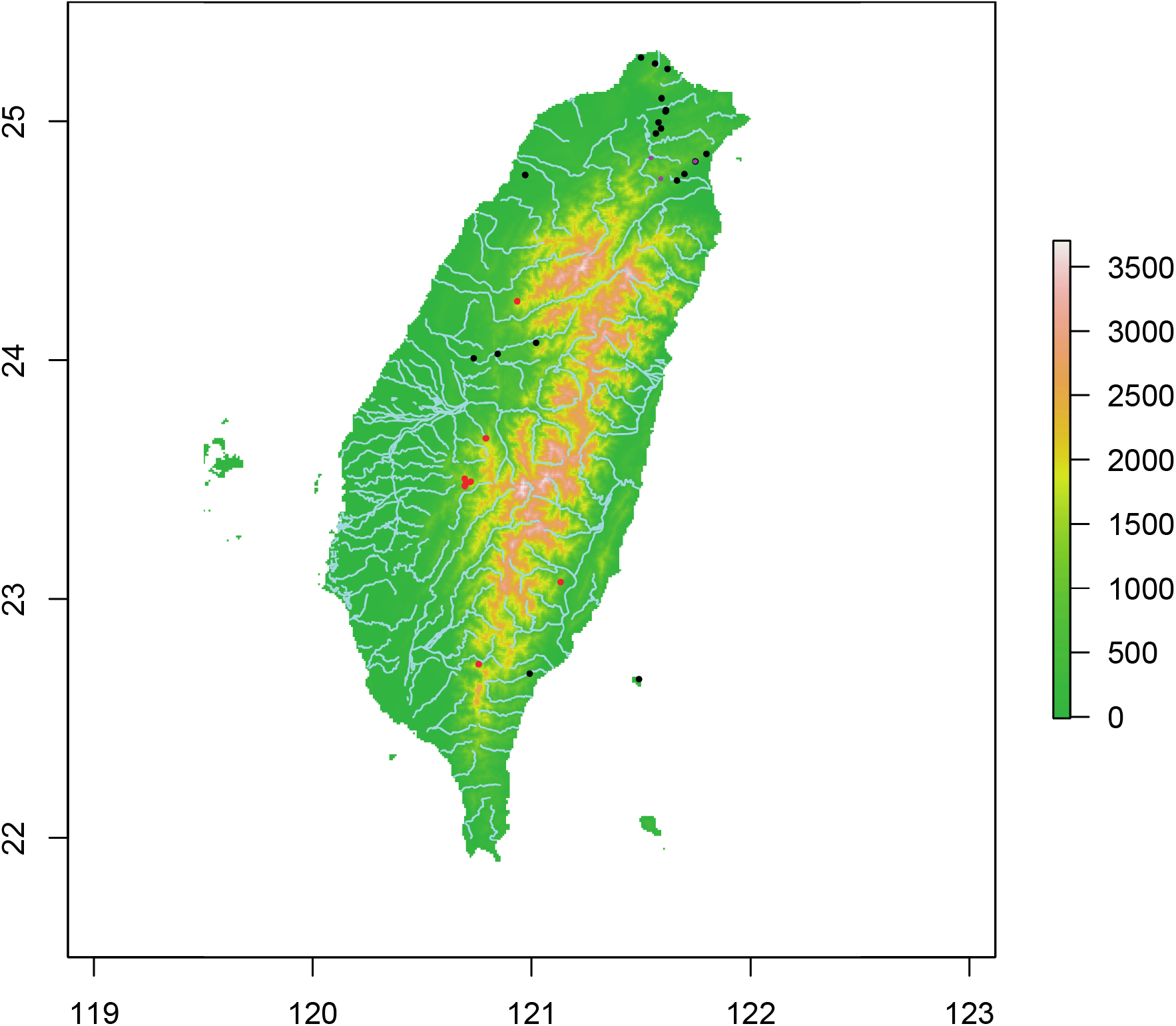
Sampling localities for *Chordodes formosanus* (black), *Acutogordius taiwanensis* (pink) and *Gordius chiashanus* (red), with a map showing elevation and river network in Taiwan. Localities from previous studies have been included. The legend shows elevation in meters.

### DNA extraction, COXI amplification and ddRADseq protocol

For DNA extraction, we used a modified version of the Qiagen DNeasy Animal Tissue Protocol (Qiagen, Hilden, Germany). We modified the original protocol by using double distilled water (ddH_2_O) instead of elution buffer for dilution, and we eluted 60 μl of DNA, given the low concentration of DNA. Additionally, we left the tissue to incubate overnight at 60°C.

The amplification of the COXI sequences through polymerase chain reaction (PCR) was done using the universal primer set LCO1490 and HC02198 (Folmer *et al.*, 1994) and was performed in a total volume of 25 μl using PuReTaq Ready-To-Go PCR Beads (Cytiva, formerly GE Healthcare, Marlborough, United States). The PCR was initiated at 95°C for 5 min, followed by 40 cycles at 95°C for 1 min, 50°C for 1 min, and 72°C for 1 min, with a final extension at 72°C for 10 min. The PCR products were purified following the NautiaZ Gel/PCR DNA Purification Mini Kit protocol (Nautia Gene, Taipei, Taiwan).

An edited version of the Peterson *et al.* (2012) protocol was used for generating the ddRADseq library. We modified the original method by using between 100 and 140 ng of genomic DNA per sample, 4 μl of ddH_2_O for bead clean-up and 60 μl of ddH_2_O for elution before measuring DNA concentration with Qubit. Single-end sequencing of 100-nucleotide base pair (bp) lengths were performed using an Illumina HiSeq2500 system.

### COXI analyses

The COXI forward and reversed sequences from each individual were visualised with MEGA X (ver. 10.1.8; Kumar *et al.*, 2018). The alignments were done with MAFFT 7.471 (Katoh and Standley, 2013) using the L-INS-i algorithm. The sequences were trimmed manually while visualising them with MEGA X.

COXI sequences at least 400 bp long for each taxon from previous studies (Chiu *et al.*, 2011, 2016, 2017, 2020) were downloaded from GenBank (Clark *et al.*, 2016: last assessed 4^th^ November 2022), aligned using MAFFT 7.471 with the L-INS-i algorithm and manually trimmed while being visualised with MEGA X. In the case of *A. taiwanensis*, a sequence of a hairworm from Myanmar (MF983649) was included for population structure analyses; said sequence has been shown to fall inside the known COXI variability of *A. taiwanensis* in previous studies (Chiu *et al.*, 2020) and therefore it should represent a sequence from said species. In total, 433 bp were recovered for 85 specimens of *C. formosanus* (36 newly generated), 414 bp for 31 *A. taiwanensis* (three newly generated) and 382 bp for 26 individual of *G. chiashanus* (one newly generated). The newly generated sequences are available in GenBank (OQ121045-OQ121047 for *A. taiwanensis*, OQ121048-OQ121083 for *C. formosanus,* OQ121084 for *G. chiashanus).*

Possible population structure was inferred by a haplotype network in PopArt (Leigh and Bryant, 2015), generated with the TCS 1.21 algorithm (Clement *et al.*, 2000). Estimation of *Ne* trends were calculated by running Coalescent Bayesian Skyline plot analyses (Drummond *et al.*, 2005) on BEAST2 (version 2.6.3: Bouckaert *et al.*, 2019). The most likely mutation model per each species was inferred using ModelTest, implemented in raxmlGUI (Edler *et al*., 2021) according to Akaike Information criterion (AIC; Akaike, 1998). A Relaxed Clock Log model (Drummond *et al.*, 2006), with a clock rate of roughly 0.0013 per million years, was set. We estimated this clock rate by aligning full COXI sequences from the only available *Chordodes* and *Gordius* mitogenomes from GenBank (MG257764 and MG257767; generated by Mikhailov *et al*. 2019), dividing the number of variable sites by the total number of sites and then dividing again by 2 before dividing by 110, which represents the estimated age in million years of the oldest known gordiid fossil (Poinar and Buckley, 2006). Note that estimates for the appearance of crown group Nematomorpha range from around the mid-Cambrian to the mid-Cretaceous (Howard *et al.*, 2022), but that these estimates include the possible divergence time of the *Nectonema* lineage from gordiids too. Therefore, we prefer to refer to the fossil datum, since it is the earliest possible divergence point for the Chordodidae/Gordiidae lineage. Additionally, the *C. formosanus* Japanese and Taiwanese samples were split for estimations, while the Myanmar sequence was removed for *A. taiwanensis*.

### ddRADseq analyses

Two different pipelines for processing the ddRAD data were used for *C. formosanus:* ipyrad (version 0.9.84: Eaton and Overcast, 2020) and Stacks (version 2.62: Rochette *et al.*, 2019). For the ipyrad pipeline, we performed de-novo assembly with a Phred Qscore offset of 43, 90% clust threshold, 0.5 maximum of heterozygotes in consensus, and minimum sample locus at 80% of all the samples, while keeping other settings at their default values. Samples with less than 300 loci were discarded for further analyses. For Stacks, the protocol listed in Rivera-Colón and Catchen (2022) was followed and all the sites (variant and invariant) were outputted in the Variant Call Format (VCF) file. In total, 27 individuals were retained for *C. formosanus* from both pipelines. In the case of *A. taiwanensis* and *G. chiashanus*, only two individuals per species had at least 200,000 reads after trimming and therefore we only used Stacks for their assemblies, with the “r” flag set at 1 instead of 0.8.

To check the genetic population structure of *C. formosanus*, tools from ipyrad and R were used with both the ipyrad and Stacks output. The Stacks VCF file was filtered to remove the loci without SNPs and the SNPs with more than 20% missing data, using an edited version of previously released R scripts (Dalapicolla *et al.*, 2021). Additionally, the filtered VCF file was converted to a HDF5 database using the converter inside the ipyrad-analysis toolkit (Eaton and Overcast, 2020) to make it compatible with the tools implemented inside ipyrad. After this, *k-means* Principal Component Analyses (PCA) were run 100 times in the ipyrad-analysis tools suite. Moreover, the *snapclust* clustering method implemented into the “adegenet” R package (Beugin et al., 2018) was used. The best number of clusters *(k)* was calculated by computing the AIC and the Bayesian information criterion (BIC; Schwarz, 1978), while picking the value for which both criteria were lower. The *snapclust* analyses were run with both the ipyrad full output STRUCTURE file and a STRUCTURE file generated from the filtered Stacks VCF thanks to PGDSpider (Lischer and Excoffier, 2012). Furthermore, for analysing possible admixture between geographically separated *C. formosanus* individuals, RADpainter and fineRADstructure (Malinsky *et al.*, 2018) were used and the results were plotted with R. The RADpainter file from the Stacks output was used for such software, while the “hapsFromVCF” command generated an input file with the ipyrad-outputted VCF.

For population demographic analyses of all the taxa, a site-frequency spectra (SFS) file was generated using the easySFS script (https://github.com/isaacovercast/easySFS) by converting the Stacks VCF file with the invariant sites, choosing the number of individuals that maximise the number of segregating sites. Note that easySFS takes into consideration the fact that missing data are present, and therefore we used the full Stacks VCF output for SFS conversion. After that, Stairway Plot 2 (version 2.1.1: Liu and Fu, 2020) was used per each species following the software’s manual. Given the current knowledge on Taiwanese horsehair worms (Chiu, 2017; Chiu *et al.*, 2020), generation time was set at 1 per year. As for the mutation rate, given the lack of whole-genome data for horsehair worms, a spontaneous mutation of rate of 2 x 10^-9^ per site per generation, which is the one of the nematode *Pristionchus pacificus* (Weller *et al.*, 2014), was set. We argue that, even if nematodes and hairworms have been split for hundreds of millions of years (Howard *et al.*, 2022), mutation rates in invertebrates tend to be around the same order of magnitude (10^-9^; e.g., Konrad *et al.*, 2019) and therefore should not influence the results in a significant way.

## Results

### COXI analyses

In total, there were 54 different mitochondrial haplotypes in *Chordodes formosanus*, 9 of which were hypothetical and 28 of which were made up by a single individual (Fig. 2A). *Acutogordius taiwanensis* presented 9 haplotypes, none of which were hypothetical and 5 of which were made up by a single individual; however, 5 haplotypes were separated from the most common one by just one mutation (Fig. 2B). 24 haplotypes, with 5 hypotheticals, were counted for *Gordius chiashanus*, and 15 of these were made up by a single individual (Fig. 2C).

**Figure 2.**
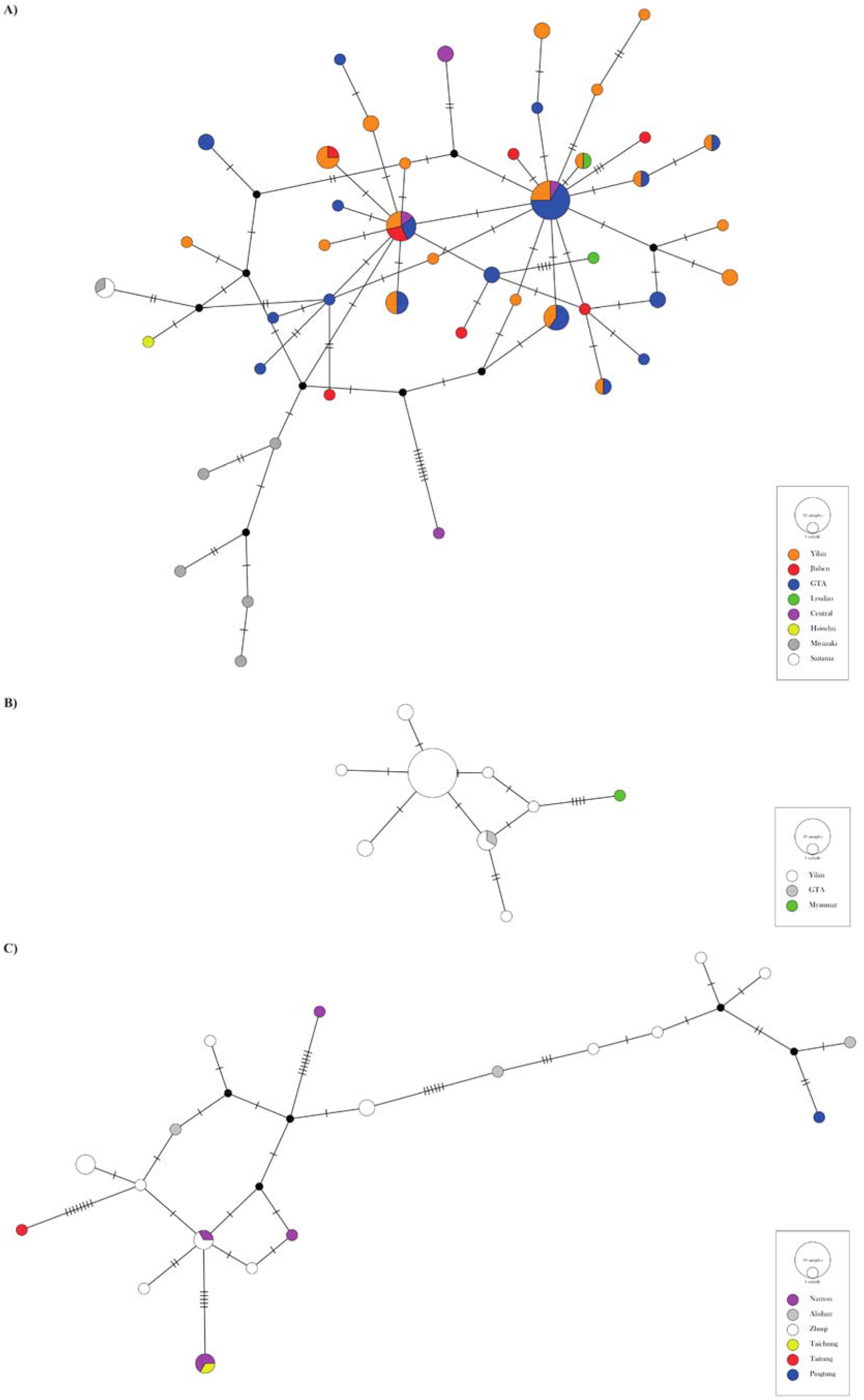
Haplotype networks based on COXI fragments. A) *Chordodes formosanus;* B) *Acutogordius taiwanensis;* C) *Gordius chiashanus.*

Nucleotide diversity (pi) was 297585 for *C. formosanus,* 0.0010493 for *A. taiwanensis* and 67063.1 for *G. chiashanus*. *C. formosanus* had 54 segregating sites and 30 parsimony-informative ones, while *A. taiwanensis* had 11 and 4 and *G. chiashanus* had 30 and 23, respectively. Tajima’s D for *C. formosanus* was 3.88852 x 10^7^, while it was −2.69148 for *A. taiwanensis* and 1.20934 x 10^7^ for *G. chiashanus;* this statistic was significant (*p*=0) for both *C. formosanus* and *G. chiashanus,* but not for *A. taiwanensis (p=* 0.999976) (Sup. Table 2).

For the Coalescent Bayesian Skyline plot analyses, the number of variable sites was 49 for the Taiwanese samples of *C. formosanus*, while it was 12 for the Japanese ones. After removing the Myanmar sequence, 7 variable sites were retained for *A. taiwanensis*. We did not change the alignment file for *G. chiashanus*, and therefore 30 variable sites remained (Sup. Data). The best substitution model for the Taiwanese samples of *C. formosanus* was TrN+G4, while TIM1 was the best for the Japanese ones. HKY+I and TPM3uf+I were the best models for *A. taiwanensis* and *G. chiashanus*, respectively (Sup. Table 3). The Bayesian Skyline Plot results show how both the populations of *C. formosanus* are having a small decline, while *A. taiwanensis’s* population tends to be stable (Fig. 3). *G. chiashanus’s* population size seemed to be increasing until recently (Fig. 3).

**Figure 3.**
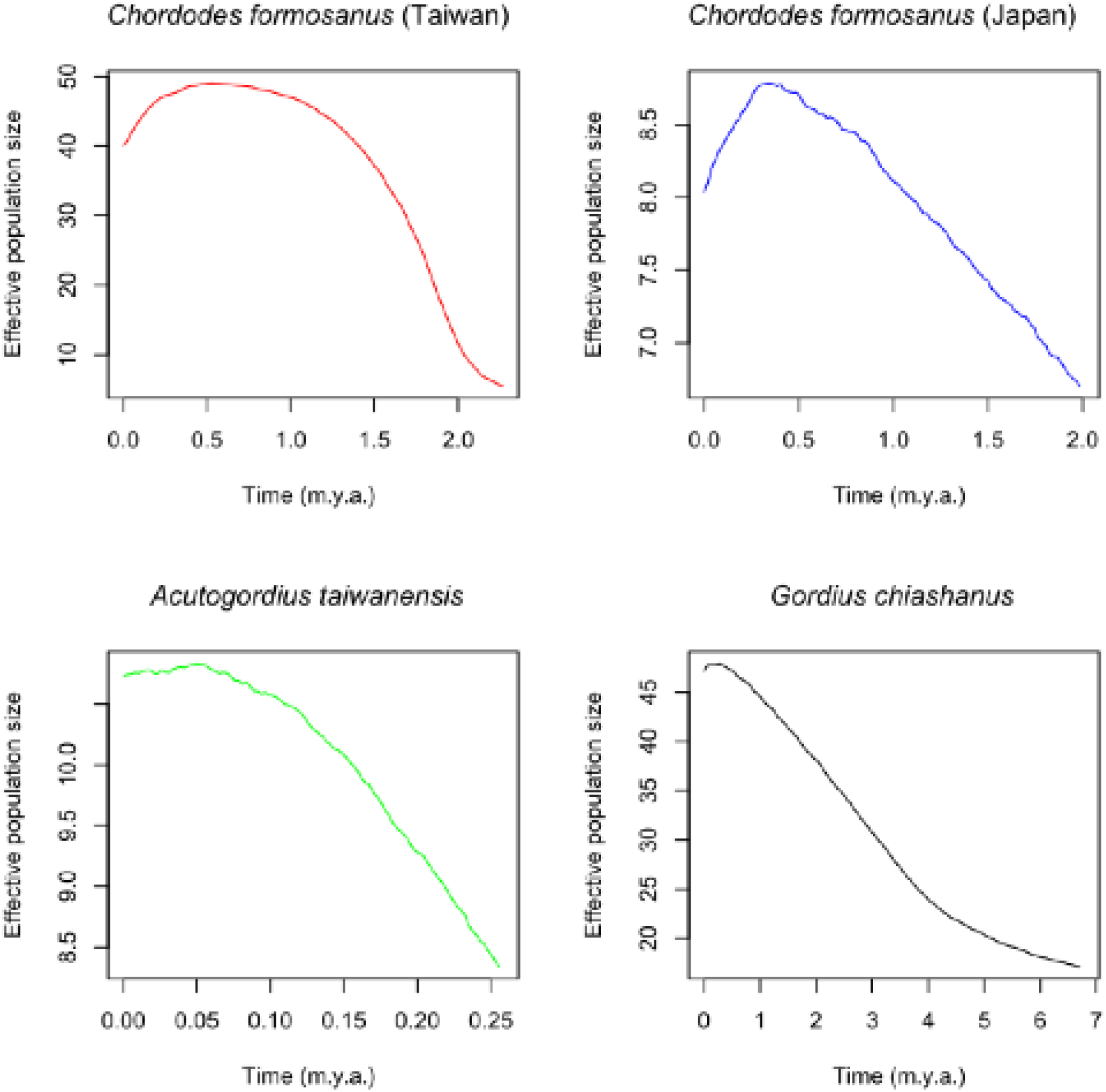
Bayesian Sky Plots based on COXI fragments per each species/population.

### ddRADseq analyses

The ipyrad trimming step led to an average read number of 2195578.39 per *C. formosanus* sample (including samples that were discarded because of the low number of reads or low number of retained loci). The ipyrad final output for *C. formosanus* had a SNP matrix size of 2248, with 17.82% missing sites, while the total sequence matrix size was 52233, with 17.59% missing sites. The lowest number of loci in assembly per an individual was 331, while the highest one was 548 and the average was 473.8888889 loci per samples (Sup. Data).

For the Stacks output, an average read number of 1672668.836 per sample was retained (including samples that were discarded because of low number of reads). The optimal settings for *C. formosanus* were obtained with both the number of allowed mismatches (M) and the number of allowed mismatches between individual loci and the catalogue of loci (n) set to 6. For *A. taiwanensis* and *G. chiashanus*, the best results were obtained with M and n set to 12 and 9, respectively. The number of R80 loci was 1876 for *C. formosanus,* 1388 for *A. taiwanensis* and 2484 for *G. chiashanus*. After removing the SNPs with 20% of the missing data, 4243 SNPs were retained for PCA and *snapclust* analyses for *C. formosanus* (Sup. Data).

Both PCA and *snapclust* analyses, conducted using both the output VCFs (ipyrad and filtered Stacks), failed to trace any structure whatsoever in *C. formosanus,* with both the AIC and BIC computed by *snapclust* agreeing that *k=1* was the optimal number of clusters for both datasets (Fig. 4; Sup. Fig. 1-2). The same pattern was observed using fineRADstructure with both input files, given that no cluster formed based on shared coancestry (Fig. 5; Sup. Fig. 3).

**Figure 4.**
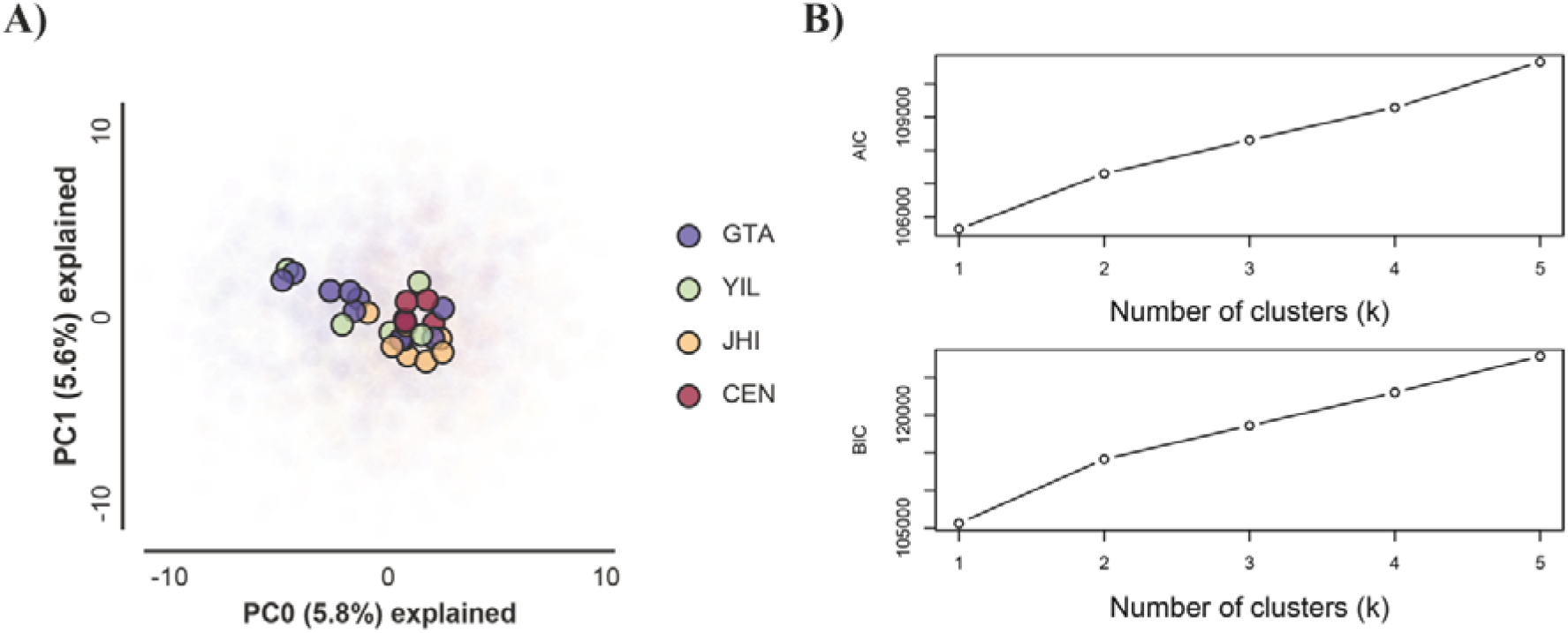
A) PCA results for Stacks ddRAD-seq data for *Chordodes formosanus*. B) AIC and BIC values from *snapclust* analyses for *C. formosanus* based on the same data as the PCA. The PCA and *snapclust* results based on ipyrad are available as Supplementary Figures 1 and 2.

**Figure 5.**
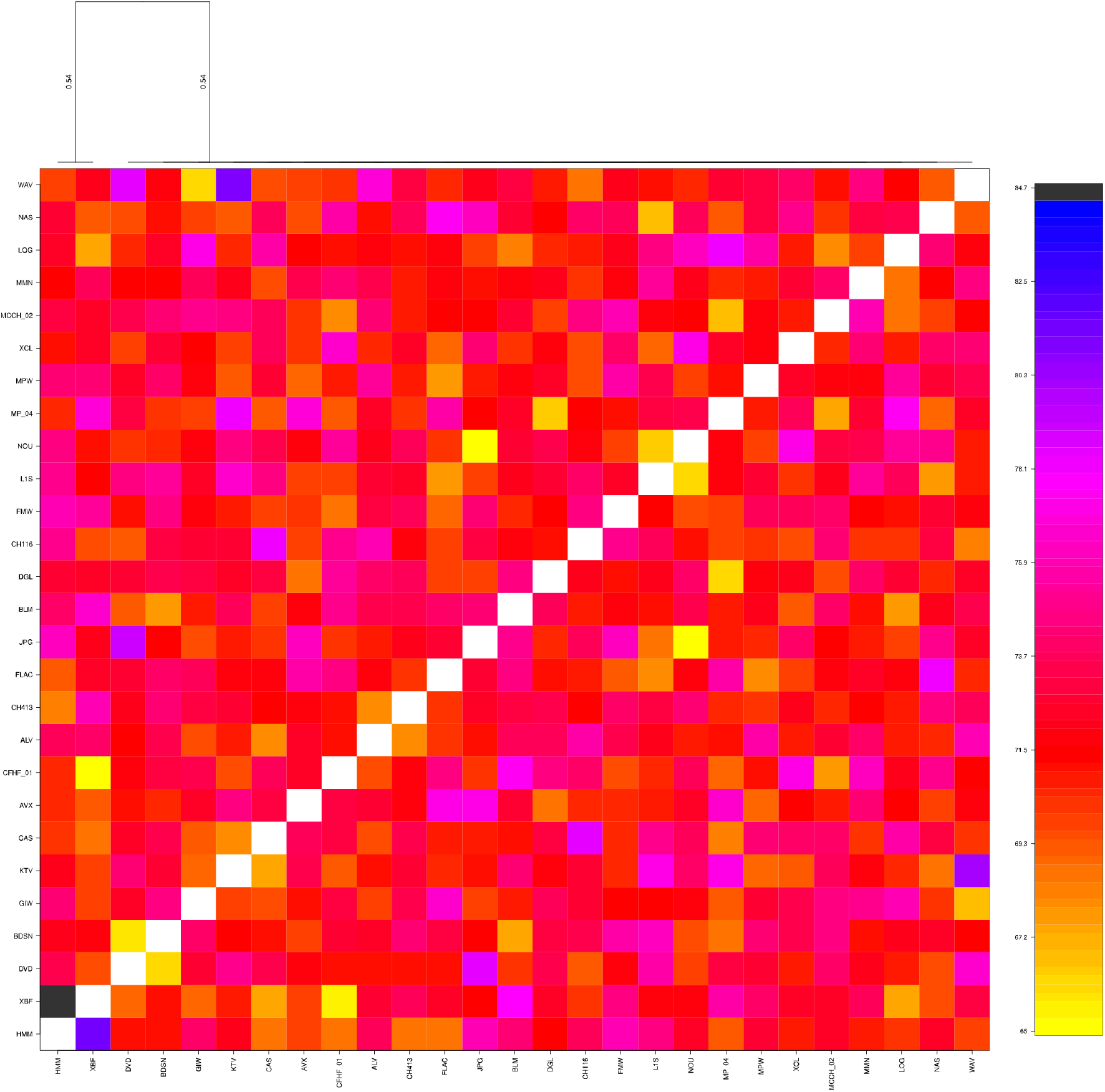
Co-ancestry matrix for *Chordodes formosanus* based on Stacks ddRAD-seq data. The one from ipyrad data is available as Supplementary Figure 3.

The Stairway Plot output for *C. formosanus* shows several population drops during the Pleistocene and an estimated *Ne* of around 1000 individuals in more recent times (Fig. 6; Sup. Fig. 4). However, only two drops were detected for both *A. taiwanensis* and *G. chiashanus,* and their estimated *Ne* in recent times were larger (around 300,000 for the former and 150,000 for the latter; Sup. Fig. 5-6).

**Figure 6.**
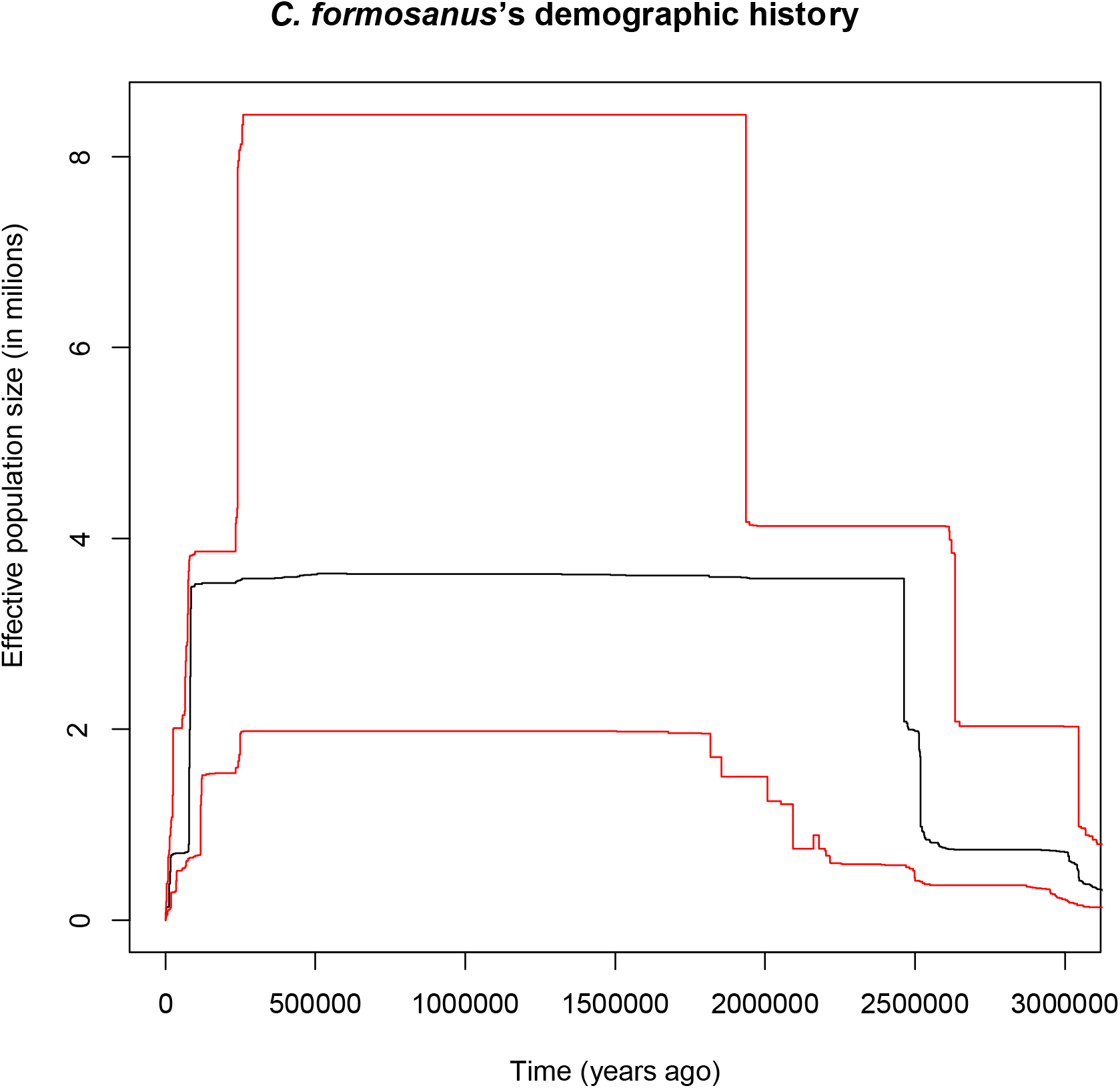
Stairway plot for *Chordodes formosanus*. The original version and the ones for *Acutogordius taiwanensis* and *Gordius chiashanus* are available as Supplementary Figures 4-6.

## Discussion

### COXI diversity analyses

Both *Chordodes formosanus* and *Gordius chiashanus* exhibit high levels of intraspecific diversity and do not exhibit a geographical genetic structure based on mitochondrial data. High mitochondrial diversity without population structure in Nematomorpha species has been shown in taxa from New Zealand (Tobias *et al.*, 2017), hinting at a high dispersal ability or a multigenerational dispersal process. In the case of the previous study, the authors were surprised to discover a lack of genetic structure in *Euchordodes nigromaculatus* (Tobias *et al.*, 2017), because the species was known to parasitise we□ta□s (family Anostostomatidae), which are definitive hosts that do not disperse well and exhibit a strong population structure (e.g., King *et al.*, 2003). Additionally, the intermediate hosts (at least two species of the stonefly genus *Stenoperla)* should not be able to disperse greatly in the mountain range inhabited by *E. nigromaculatus* (Tobias *et al.*, 2017). However, new studies have revealed that this New Zealand species can also infect the cockroach *Celatoblatta quinquemaculata* (Doherty *et al.*, 2022), a common insect in the mountains of central Otago (Worland *et al.*, 2004), which is the area considered by Tobias *et al.* (2017). This suggests that this cockroach shows more dispersal ability than the we□ta□s. Therefore, it could be the case that dispersal is facilitated by definitive hosts. Note that dispersal by definitive hosts does not exclude the possibility of a multigenerational process, as highlighted by Tobias *et al.* (2017). Given the scarce knowledge of intermediate hosts used by Taiwanese horsehair worms, we should not exclude them as possible vectors too.

In the case of *G. chiashanus,* only one definitive host (a millipede species from the genus *Spirobolus)* and only one potential intermediate host (the mayfly *Ephemera orientalis)* [Chiu *et al.*, 2020], which is known to be present in different East Asian nations [Lee *et al.*, 2008]) are known. The mayfly does not seem to be commonly infected (Chiu *et al.*, 2020), which raises questions about both the intermediate and definitive host ranges of *G. chiashanus.* Another species with similar ecology, *Gordius terrestris,* prefers earthworms as natural intermediate hosts (Anaya *et al.*, 2021) and *G. chiashanus* might do the same. However, these considerations do not change the fact that *G. chiashanus* can be found in different mountain localities (Chiu *et al.*, 2020; this study), and its COXI haplotypes suggest a high dispersal ability that allows it to colonize such mountain areas. Furthermore, all the possible explanations for dispersal listed here (by paratenic hosts, by definitive hosts, by multigenerational process) do not exclude each other, and it is likely that hairworm dispersal is a complex process that may include all the options listed above.

While *Acutogordius taiwanensis* showed less intraspecific genetic diversity than the other two considered taxa, it must be noted that almost all the samples for this species came from a single region of Taiwan, Yilan County (Chiu *et al.*, 2017; this study). Meanwhile, the COXI sequences of *C. formosanus* were generated from individuals from multiple localities in Taiwan and Japan (Chiu *et al.*, 2011, 2016, 2017; this study), and the ones for *G. chiashanus* came from multiple middle elevation mountain localities in central-south Taiwan (Chiu *et al*., 2020; this study). These differences in sampling areas can be explained by the varying difficulties encountered when sampling the adult horsehair worms, which are usually regarded as significant issues in studying them (Schmidt-Rhaesa, 2012; Bolek *et al.*, 2015). *C. formosanus* is mostly a diurnal species, like the mantises it usually parasitises (Chiu *et al*., 2011; Chiu, 2017), and can often be found road-killed with its hosts (Chiu, 2017). In the case of *G. chiashanus*, although sampling the adult with the definitive hosts can be challenging (Chiu, 2017), free-living adults tend to aggregate together during breeding season, which allows for the collection of several individuals at the same time (Chiu *et al.*, 2020). *A. taiwanensis*, however, is mostly a nocturnal species because of the habits of its definitive hosts; in fact, it has been recorded in two different raspy cricket species (family Gryllacrididae) and 9 different katydid taxa (family Tettigoniidae; Chiu *et al.*, 2017), which are mostly nocturnal animals (Chiu, 2017). Both host families can be hard to sample because of their biology. More precisely, raspy crickets tend to burrow inside the soil and have an inconspicuous brown colouration (Rentz and John, 1990), while katydids are known for their leaf-like external morphology (Mugleston *et al.*, 2016). Additionally, the infection rate and the population size seem to be lower in *A. taiwanensis* than in *C. formosanus*, which roughly occupy the same environments (M. C. Chiu, pers. comm.). As a result, it is highly likely that the genetic diversity of *A. taiwanensis* is extremely underestimated, as already suggested by Chiu (2017) and Chiu *et al*. (2020), and the species may have gone unnoticed in many areas, as has happened with other parasites (Pappalardo *et al.*, 2020). On the other hand, *A. taiwanensis* can be the most vulnerable Taiwanese hairworm species to human intervention (Chiu *et al.*, 2016); previous studies highlighted the possibility of hairworm extirpation caused by anthropic activities (Sato *et al.*, 2014). Therefore, *A. taiwanensis* may have gone extinct in some areas in Taiwan due to human intervention, especially in areas where agricultural land expanded in the past or in recent years (Chen *et al.*, 2019). This potential extirpation may also explain the lack of *C. formosanus* reports in highly farmed areas (De Vivo and Huang, 2022) and highlight the possible reaction of hairworms to pesticides (Achiorno *et al*., 2018).

### *ddRADseq diversity analyses for* Chordodes formosanus *in Taiwan*

Even with genome-wide data, *C. formosanus* did not show any sign of population structure according to geographical origins (Fig. 4). Together with its known distribution and COXI haplotype, the results suggest a panmictic population with great dispersal ability. However, gordiids disperse poorly by themselves when they are at the larval or adult stage (Schmidt-Rhaesa, 2012; Bolek *et al.*, 2015; Chiu *et al.*, 2016; Chiu, 2017) and the river network in Taiwan prevents non-flying organisms associated with freshwater from dispersing (Shih *et al*., 2006). Therefore, dispersal by host (intermediate and/or final ones) with or without a multigenerational process can be a possible explanation for this pattern. Previous modelling efforts noticed a very high similarity between the mantis hosts’ range and the range of *C. formosanus*, hinting at a potential dispersal by definitive hosts (De Vivo and Huang, 2022). Currently, the intermediate hosts of *C. formosanus* are only partially known (Chiu, 2017) and include non-biting midges in the family Chironomidae, the caddisfly *Chimarra formosana* and stoneflies in the genus *Kamimuria* (Chiu *et al.*, 2016; Chiu, 2017), with a seeming preference for the dipteran family (Chiu *et al.*, 2016). There has been no study on the dispersal ability of said clades in Taiwan, but it is known that the dispersal abilities in midges, stoneflies and caddisflies vary greatly among taxa (Ferrington, 2008; Arce *et al*., 2021). Nevertheless, the preference for active dispersing flying insects as intermediate and definitive hosts can explain the lack of geographic structure shown in our results.

### Demographic history inferences

The demographic analyses using the COXI datasets were able to trace historical demography through 2 million years for *C. formosanus* (with the Taiwanese population going a little bit deeper in time). The same approach revealed *Ne* trends up to roughly 6 million years ago for *Gordius chiashanus* and 250,000 years of demographic history for *Acutogordius taiwanensis*. This difference in scale may have been caused by the lower variability of the *A. taiwanensis* sequences (7 variable sites in total), which in turn was caused by a limited geographic sampling. In general, it is known that mitochondrial sequence data usually tend to recover demographic histories in the Pleistocene (e.g., Huang and Lin, 2010); however, such data is less informative with recent history (Nunziata and Weisrock, 2018). The COXI-based Bayesian Skyline Plots revealed recent drops in *Ne* for both Taiwanese and Japanese populations of *C. formosanus* over the last 250,000 years. Several drops in the same period were found by Stairway Plot using the ddRADseq data. Additionally, both approaches found a demographic increase a little bit before 2 million years ago for the Taiwanese population. That said, Stairway Plot analyses with ddRADseq data were able to detect more events in the last 100,000 years for the Taiwanese population of *C. formosanus*, compared to the Coalescent Bayesian Sky Plot based on COXI sequences.

The demographic histories reconstructed using genome-wide SNP data and the Stairway Plot were very large for *A. taiwanensis* and *G. chiashanus*, but at the same time, the software failed to recover any possible trends in recent years. This was caused by the very small sample size, which was 2 diploid individuals per each species. Specifically, the number of historical events that can be estimated using Stairway Plot, and any programs that take site frequency spectrum as inputs and calculate composite likelihoods, is constrained to the number of site frequency categories. As a result, for a folded site frequency spectrum, as implemented here, the use of two individuals will only result in two site frequency categories, and thus only two historical demographic episodes can be estimated (Liu and Fu, 2020). The two species also have extremely large *Ne* estimated based on the Stairway Plot results. It is known that sample size can impact the estimated demographic parameters using RADseq data (Nunziata and Weisrock, 2018) and therefore we attribute the high *Ne* estimates to the sampling size.

For *C. formosanus,* however, way more events were recorded from the Stairway Plot analyses. That said, it is surprising that this species, which is regarded as the most common in Taiwan, is showing signs of decline—although the current *Ne* should still be way larger than enough to sustain the population (Franklin, 1980; Pérez□Pereira *et al.*, 2022). The recent demographic decrease might be the result of extirpation events (Legendre *et al.*, 2008) possibly caused by changes in land use in Taiwan (De Vivo and Huang, 2022), given that hairworms are sensitive to human activities and pollution (Poinar, 2008; Sato *et al.*, 2014; Chiu *et al.*, 2016; Achiorno *et al.*, 2018).

## Conclusions

In this study, we evaluated the use of different genetic and analytical tools for estimating demographic history and population genetic structure. In our datasets, sampling bias for some taxa was present and can influence the results. For possible future studies on the conservation genetics of parasites, we give three suggestions that can help with evaluating a parasite’s conservation status:

i. consider the target species’ ecologies, given that they can highly influence their population structure (Van Schaik *et al*., 2015; Radac□ovská *et al.*, 2022), *Ne* estimates (Criscione and Blouin, 2005; Criscione *et al.*, 2005; Criscione, 2013, 2016; Strobel *et al.*, 2019; Doña and Johnson, 2020) as well as the sampling strategy, given that some parasites have life cycles that can influence when and how to collect them (e.g., Van Schaik *et al.*, 2015);
ii. try to get the geographically broadest and biggest sample size possible; for example, in this study a small sample size in the ddRADseq dataset did not allow us to estimate enough recent trends for two taxa, while the COXI dataset for *A. taiwanensis* was too biased to one area, which did not allow us to properly estimate the diversity of the species;
iii. define the molecular and bioinformatic tools used; a direct estimate of *Ne* for parasites can be tricky to calculate (Criscione, 2013; Strobel *et al.*, 2019; Carlson *et al.*, 2020). Given this, we suggest focusing on trends instead of raw numbers. Additionally, previous studies showed possible limits of single-locus analyses (Vitalis and Couvet, 2001; Ho and Shapiro, 2011). Therefore, we suggest using as many loci as possible. An approach such as RAD or ddRADseq can be used for such a goal, given that they provide an approximation of the genome. However, single-locus data are often the kinds of data that are the most available for parasites (Selbach *et al.*, 2019). Therefore, if budget and resources are limited, Sanger sequencing can be pursued (but see Radaco□oská *et al.* 2022 for caveats with mitochondrial sequences in polyploid species). The use of depositories such as GenBank can be useful for retrieving previously released sequences and therefore increase both sample size and loci used while utilising software such as BEAST2 that can use such data. That said, it is crucial to check for a possible population structure before demographic trend analyses, since it influences the results (Ho and Shapiro, 2011).

## Supporting information

Supplementary Images

## Data

The COXI sequences produced in this study are available in GenBank (accession numbers: OQ121045-OQ121084). The ddRAD Stacks-trimmed reads are available in Sequence Read Archive (BioProject number: PRJNA914055). Supplementary Data (i.e., logs and outputs) for this article can be found in Zenodo (https://zenodo.org/record/7659651, doi: 10.5281/zenodo.7659651).

## Acknowledgements

We would like to thank all the people who helped us collect samples all around Taiwan. We would also like to thank Ming-Chung Chiu for helping us with designing the sampling strategy and discussing the paper and the biology of the involved species, Hsiang-Yun Lin for sampling assistance, Justin Pelofsky for revising the grammar of this article, Xiaoming Liu for helping us with Stairway Plot 2’s settings, the DNA Sequencing Core Facility of the Institute of Biomedical Sciences, Academia Sinica for Sanger sequencing and the NGS High Throughput Genomics Core at Biodiversity Research Center, Academia Sinica, for providing Illumina sequencing.

## Author Contributions

MDV extensively sampled the specimens around the country, performed the bioinformatic analyses and wrote the original draft. MDV and WYC performed molecular work. MDV and JPH conceived and designed the study. All the authors designed the methodology, corrected the original draft and approved the manuscript’s content.

## Ethical Standards

Not applicable.

