## Supplementary Images for "Testing the efficacy of different molecular tools for parasite conservation genetics: a case study using horsehair worms (Phylum Nematomorpha)"

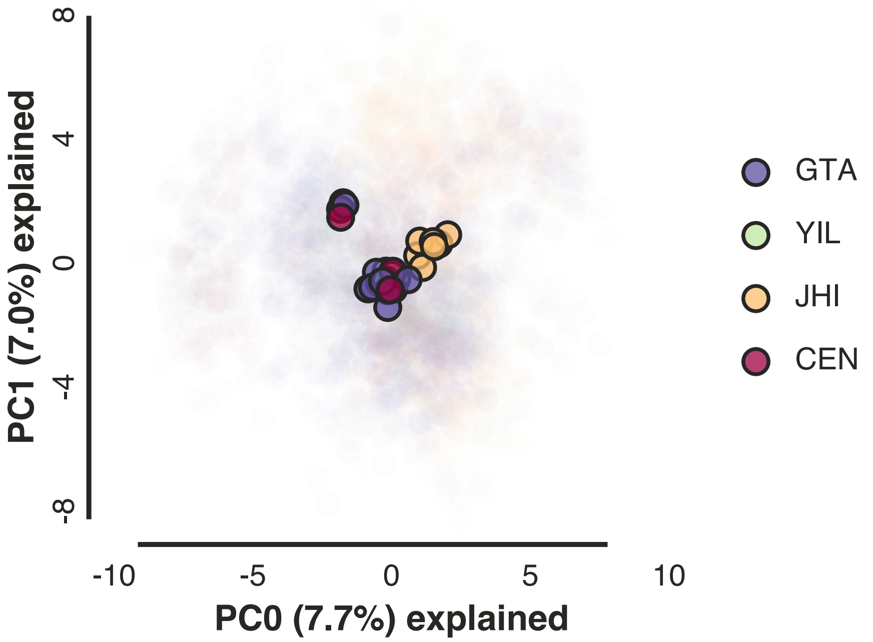


**Supplementary Figure 1**. PCA results for ipyrad ddRAD-seq output data for *Chordodes formosanus*.

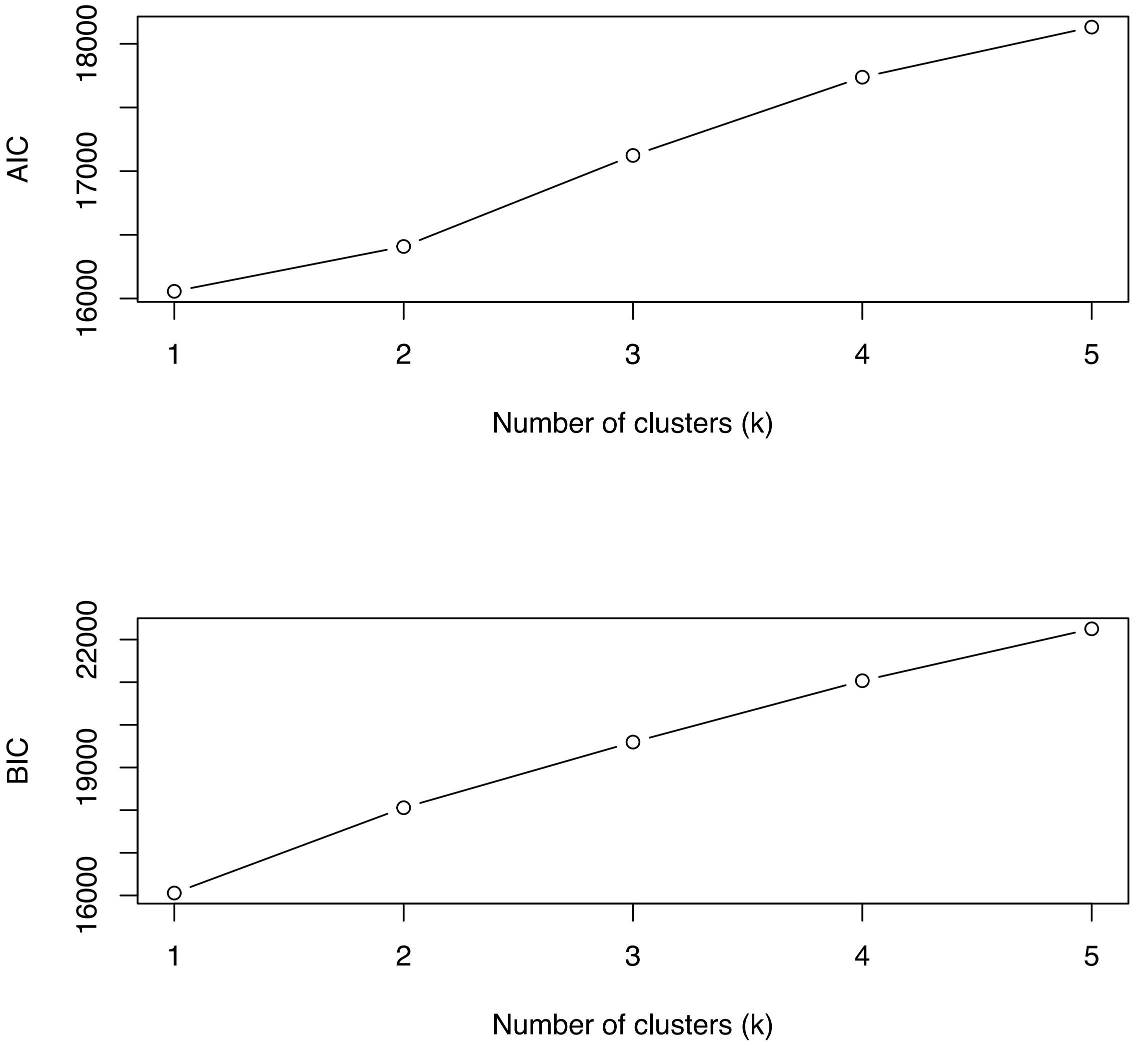


**Supplementary Figure 2**. AIC and BIC values from *snapclust* analyses for ipyrad ddRAD-seq output data for *Chordodes formosanus*.

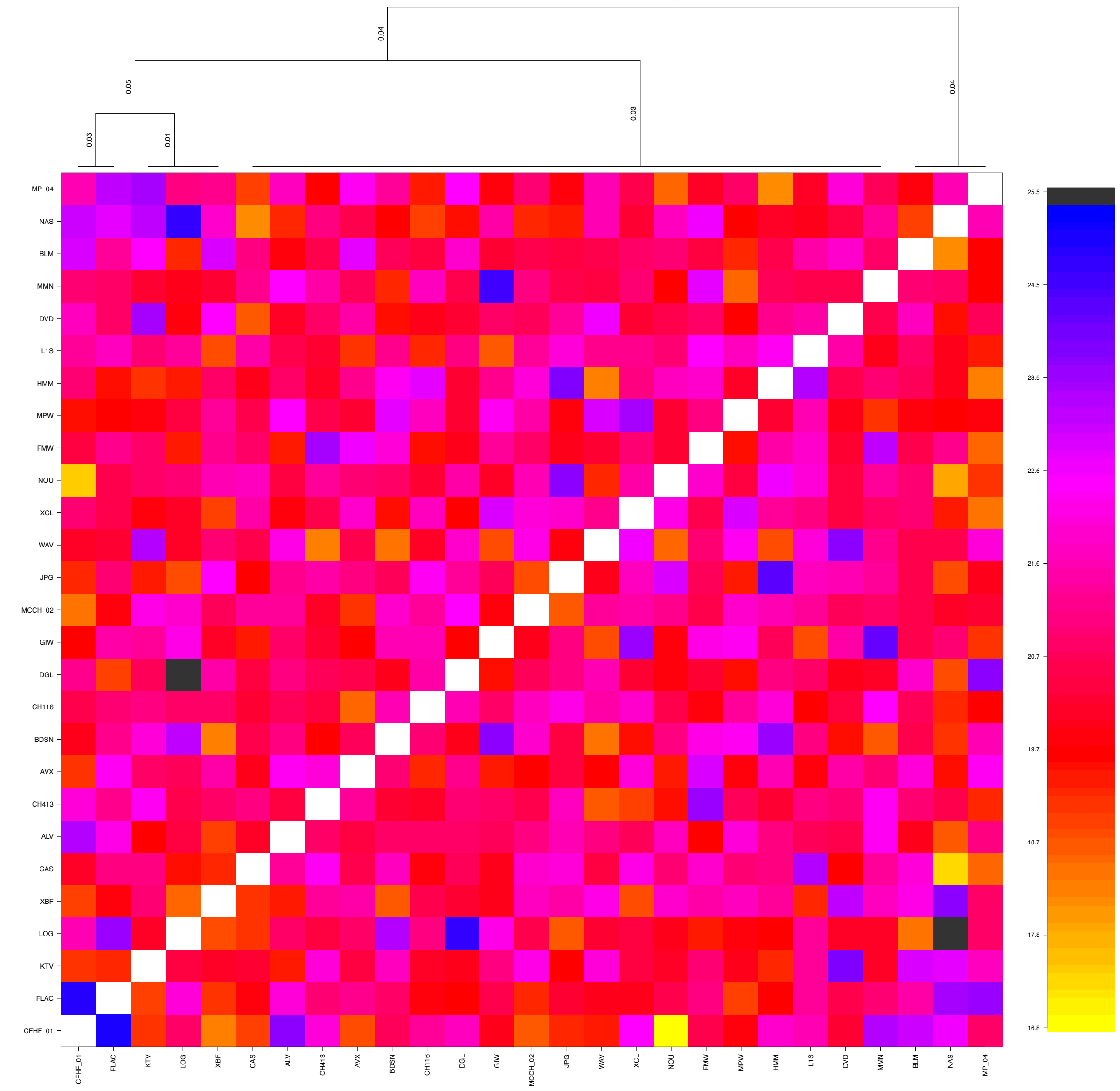


**Supplementary Figure 3**. Co-ancestry matrix for *Chordodes formosanus* based on ipyrad ddRAD-seq output data.

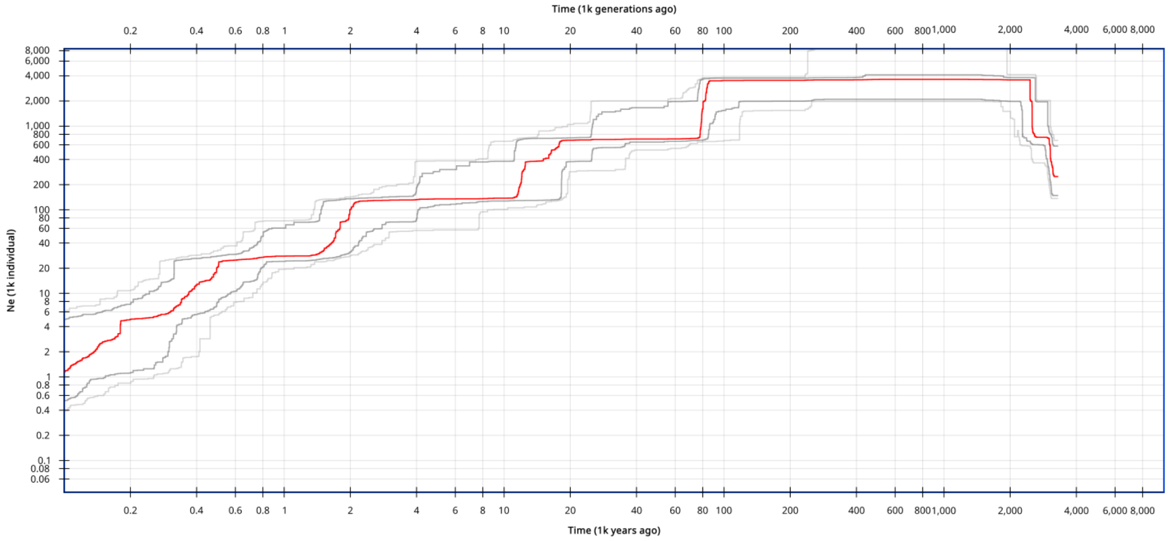


**Supplementary Figure 4**. Original Stairway plot for *Chordodes formosanus*.

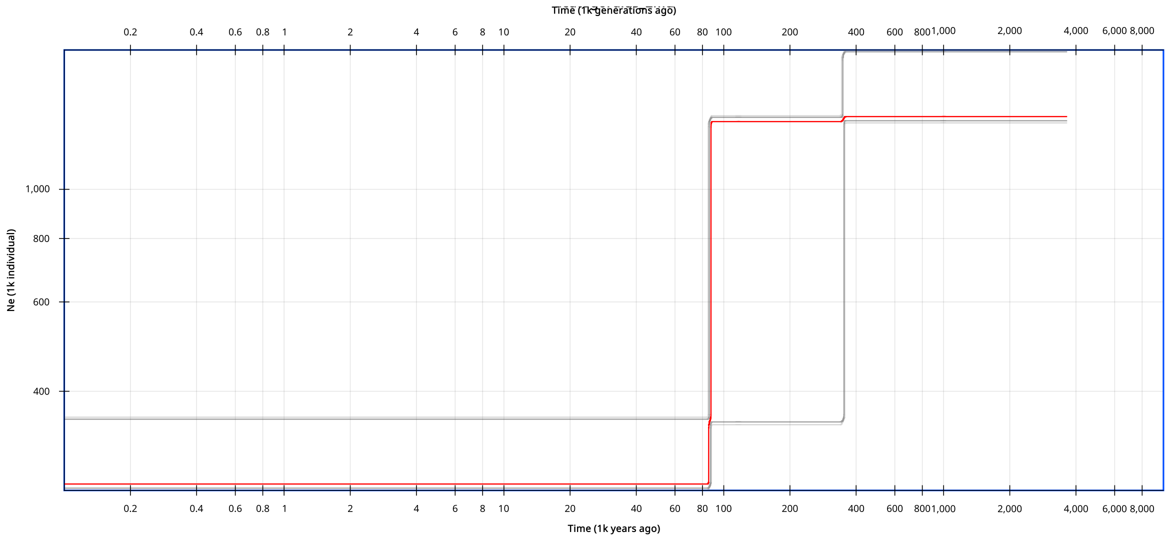


**Supplementary Figure 5**. Stairway plot for *Acutogordius taiwanensis*.

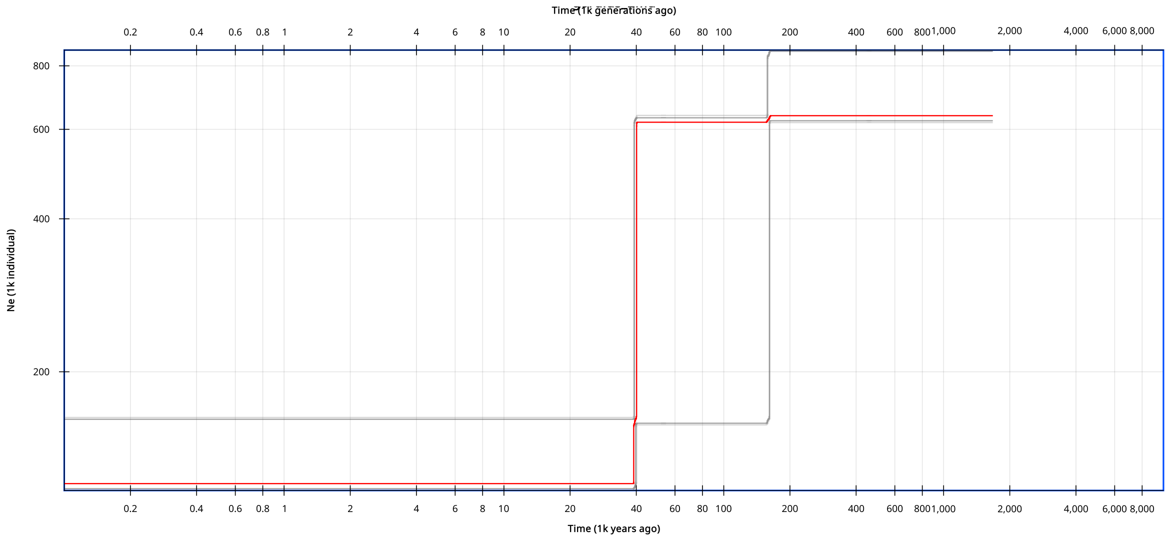


**Supplementary Figure 6**. Stairway plot for *Gordius chiashanus*.
